# A robust association test leveraging unknown genetic interactions: Application to cystic fibrosis lung disease

**DOI:** 10.1101/2023.06.22.546041

**Authors:** Sangook Kim, Lisa J Strug

**Affiliations:** Biostatistics Division, Dalla Lana School of Public Health, University of Toronto, Toronto, ON, Canada; Genetics and Genome Biology, The Hospital for Sick Children, Toronto, ON, Canada; Department of Statistical Sciences, University of Toronto, Toronto, ON, Canada; Department of Computer Science, University of Toronto, Toronto, ON, Canada; The Centre for Applied Genomics, The Hospital for Sick Children, Toronto, Ontario, Canada

## Abstract

For complex traits such as lung disease in Cystic Fibrosis (CF), Gene x Gene or Gene x Environment interactions can impact disease severity but these remain largely unknown. Unaccounted-for genetic interactions introduce a distributional shift in the quantitative trait across the genotypic groups. Joint location and scale tests, or full distributional differences across genotype groups can account for unknown genetic interactions and increase power for gene identification compared with the conventional association test. Here we propose a new joint location and scale test (JLS), a quantile regression-basd JLS (qJLS), that addresses previous limitations. Specifically, qJLS is free of distributional assumptions, thus applies to non-Gaussian traits; is as powerful as the existing JLS tests under Gaussian traits; and is computationally efficient for genome-wide association studies (GWAS). Our simulation studies, which model unknown genetic interactions, demonstrate that qJLS is robust to skewed and heavy-tailed error distributions and is as powerful as other JLS tests in the literature under normality. Without any unknown genetic interaction, qJLS shows a large increase in power with non-Gaussian traits over conventional association tests and is slightly less powerful under normality. We apply the qJLS method to the Canadian CF Gene Modifier Study (n=1,997) and identified a genome-wide significant variant, rs9513900 on chromosome 13, that had not previously been reported to contribute to CF lung disease. qJLS provides a powerful alternative to conventional genetic association tests, where interactions my contribute to a quantitative trait.

**Author summary:** Cystic fibrosis (CF) is a genetic disorder caused by loss-of-function variants in CF transmembrane conductance regulator (*CFTR*) gene, leading to disease in several organs and notably the lungs. Even among those who share identical CF causing variants, their lung disease severity is variable, which is presumed to be caused in part by other genes besides *CFTR* referred to as modifier genes. Several genome-wide association studies of CF lung disease have identified associated loci but these account for only a small fraction of the total CF lung disease heritability. This may be due to other environmental factors such as infections, smoke exposure, socioeconomic status, treatment of lung diseases or a numerous other unknown or unmeasured factors that may interact with modifier genes. A class of new statistical methods can leverage these unknown interactions to better detect putative genetic loci. We provide a comprehensive simulation study that incorporates unknown interactions and we show that these statistical methods perform better than conventional approaches at identifying contributing genetic loci when the assumptions for these approaches are met. We then develop an approach that is robust to the typical normal assumptions, provide software for implementation and we apply it to the Canadian CF Gene Modifier Study to identify novel variants contributing to CF lung disease.

## Introduction

Conventional association tests in genome-wide association studies (GWAS) aims to detect a change in the conditional mean for a quantitative phenotype across genotypic groups at a genetic polymorphism. For a bi-allelic single nucleotide polymorphism (SNP), Paŕe et al. (2010) demonstrated that an unknown interaction of a gene with another (G x G) or with an external environmental factor (G x E) can induce a change in the conditional distribution of the phenotype across the genotypic groups, which manifests as heterogeneity in the conditional variance. Hence, an unknown genetic interaction can be detected through a variance test. Since genetic interactions are generally unknown or difficult to measure *a priori*, this indirect approach, which does not require specification of variables interacting with a given polymorphism, is a convenient way to detect contributing genetic variants that may be obscured by unknown genetic interactions. As a result, there has been a renewed interest in the classical tests for variance heterogeneity *e.g.* Bartlett’s test (Bartlett, 1937) and Levene’s test (Levene, 1961). These tests were originally developed to verify the underlying assumption in the analysis of variance. More recently, a class of new statistical methods test for heterogeneity in both the conditional mean and variance jointly or even in higher moments or in the phenotypic distribution across the genotypic groups. Through simulation studies, Soave et al. (2015) showed that, in the presence of an unknown genetic interaction, these methods can be more powerful than the conventional location test. The improvement in power is especially important for GWAS of rarer diseases such as Cystic Fibrosis (CF) where the total number of cases available to study is limited.

Several joint location and scale tests have been developed (Cao et al., 2014; Rönnegård and Valdar, 2011; Soave et al., 2015; Staley et al., 2021), and they generally jointly model the location and scale of the conditional phenotypic distribution, assuming the errors are normally distributed. The main difference between these joint location and scale tests lies in the estimation of parameters for the conditional variance and the iterative process to fit the joint models. The parameters for the conditional variance are estimated either based on squared residuals or absolute deviations, analogous to Bartlett’s test and the Brown-Forsythe test (Brown and Forsythe, 1974), respectively, for discrete predictors. In joint location and scale tests such as the joint location-scale test (Soave et al., 2015; Soave and Sun, 2017) and joint location-scale score test (Staley et al., 2021), the location and scale parameters are fitted separately whereas, in others such as the likelihood ratio test (Cao et al., 2014) and double generalized linear model (Rönnegård and Valdar, 2011), the iterative process cycles between the location model and the scale model, allowing for joint estimation of the location and scale parameters. Although these methods perform well under normality, phenotypes are commonly non-normal. Rank-based inverse normal transformation is a popular data transformation technique in genetic epidemiology, as a remedial measure for non-normal error (Beasley et al., 2009; McCaw et al., 2020; Rönnegård and Valdar, 2011; Soave et al., 2015). However, the impact of applying this transformation to the joint location and scale tests on the type I error and power is not well understood. We provide a review of these joint location and scale tests and conduct simulation studies to assess the impact of applying rank-based inverse normal transformations on the type I error and power, under non-normality. We then propose an alternative.

Another category of methods aims at detecting changes in the mean, variance and beyond (Hong et al., 2017) or in the unconditional quantiles (Aschard et al., 2013) without requiring any distributional assumption. Although these methods are robust to non-normal error, they tend to fare much worse than the joint location and scale tests that assume normality when the distribution of the error term is close to a normal distribution (Soave et al., 2015). Additionally, these methods are not designed to incorporate continuous covariates, which can lead to spurious results if these confound the genetic association. Lastly, computational efficiency makes these methods difficult to implement genome-wide.

Quantile regression (Koenker and Bassett, 1978) provides a natural and robust framework to study the conditional distribution without making any distributional assumptions. Its applications in genomics have been growing in recent years (Briollais & Durrieu, 2014). Miao et al. (2022) developed a variance test based on quantile regression to detect an unknown genetic interaction. Here, we propose a new joint location and scale test based on quantile regression that does not require a normality assumption. Our method is robust to non-normal error and is computationally efficient, while its power under normality is comparable to other joint location and scale tests that assume normality. We compare the performance of these tests to our new test in simulation studies that include skewed and heavy-tailed error distributions. Finally, we apply our method in a GWAS of lung disease in Canadians with CF.

## Methods

For illustrative purposes, we consider a simple scenario with a single binary exposure variable interacting with a bi-allelic SNP. It can trivially be generalized to scenarios with multiple exposure variables of continuous or discrete types, or other polymorphisms. We note that the exposure does not necessarily have to be environmental (*i.e.* G x E interaction) and can involve another SNP (*i.e.* G x G interaction). However, we caution that the interacting variables must be independent from the SNP of interest since any dependence will confound the association (Dudbridge and Fletcher, 2014).

### Notation and genetic interaction model

Let *Y_i_* denote a quantitative trait, *G_i_*, the number of minor alleles, *E_i_*, a single binary exposure status and ***Z****_i_*, a vector of *p* covariates for the *i^th^* individual in a sample of size *N* . We use the bold character to indicate a matrix. For simplicity, we suppress the subscript *i* whenever it is convenient. The data generating model with an interaction between *G* and *E* is specified as follows:

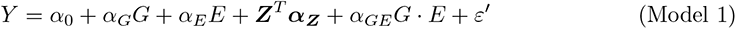

where *α*_0_*, α_G_, α_E_,* ***α_Z_*** and *α_GE_* are respectively the intercept, effect of *G*, effect of *E*, effects of ***Z*** and the interaction between *G* and *E*, and *ε*′ is an independent and identically distributed (IID) random error with a mean 0 and variance *σ*^2^. In practice, the exposure *E* is generally unknown and/or unmeasured *a priori*. Therefore, the conventional association test ignores this genetic interaction and relies on the following working model:

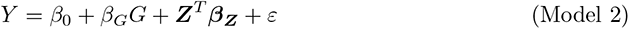

where *β*_0_*, β_G_* and ***β_Z_*** are the new intercept, effect of *G* and effects of ***Z***, respectively, and *ε* is the new error, assumed to be IID. However, in the presence of an unknown genetic interaction (*i.e. α_GE_* ≠ 0), the new error *ε* is no longer identically distributed. We illustrate the effect of the unknown genetic interaction in Figure 1 by simulating data from Model 1 based on a standard normal error.

**Figure 1:**
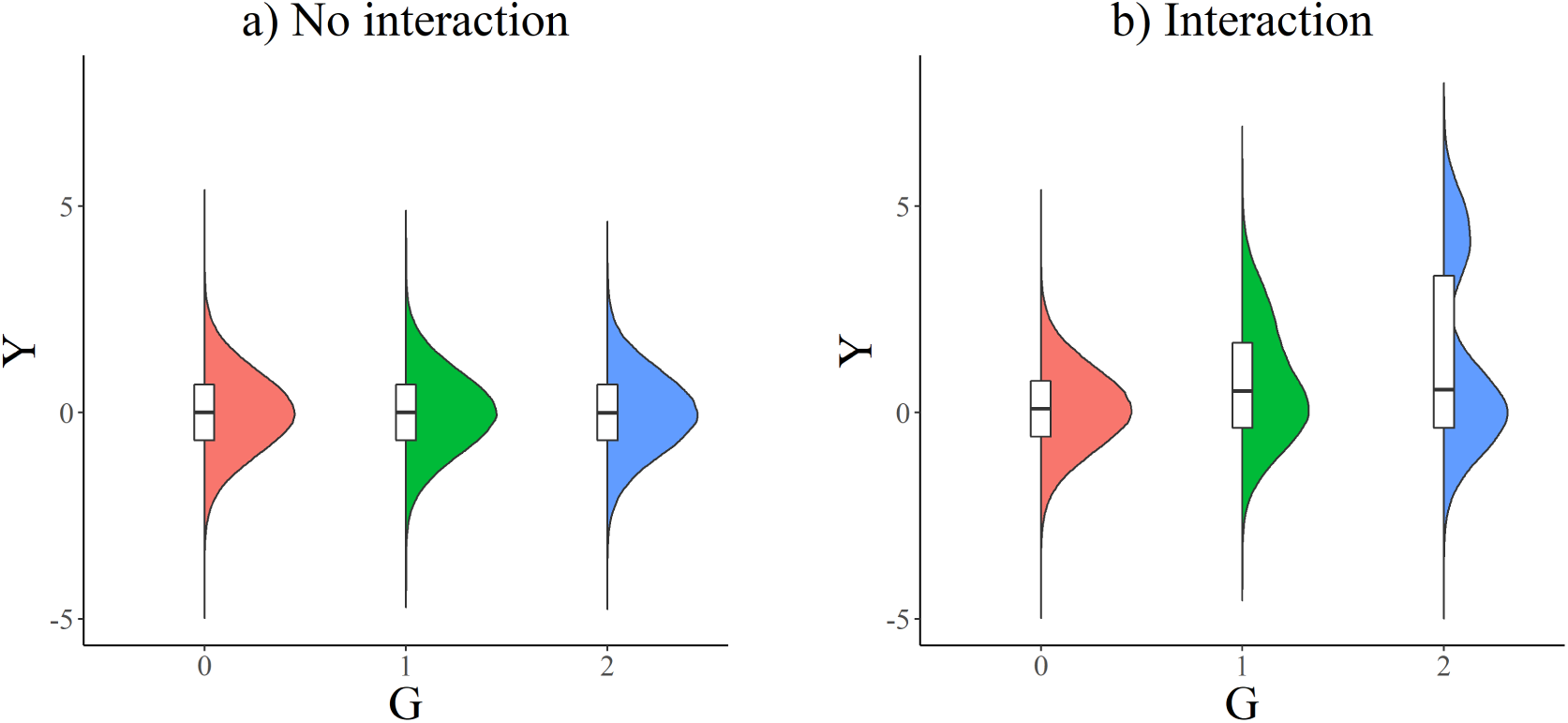
Illustration of the effect of a binary variable *E* interacting with a bi-allelic variant *G* on a Gaussian phenotype. ***Y*:** the data were simulated based on Model 1 with no covariates ***Z***, fixing the main effects *β_G_* = 0 and *β_E_* = 0.3, and interaction effect *β_GE_* = 2 with a sample size of 1,000,000. *E* was generated from a Bernoulli random variable with a probability of success *p_E_* = 0.3 and *G* was generated based on a minor allele frequency *MAF* = 0.3.

When the genetic interaction is not accounted for, the shape of the phenotypic distributions are shifted across the genotypic groups. In this particular simulation in which the exposure *E* was binary, the phenotypic distributions are pulled further apart from the unexposed to the exposed group when the number of minor alleles increases, resulting in a bi-modal shape.

Based on this finding, the unknown genetic interaction can be detected by measuring the shift in the phenotypic distributions across the genotypic groups (Aschard et al., 2013). Since the shift in the phenotypic distributions induces variance heterogeneity, Paŕe et al. (2010) proposed to capture the unknown genetic interaction by a variance test. They provided an explicit form of the conditional phenotypic variance for a given genotypic group as follows:

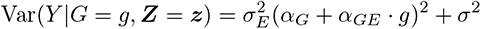

where *σ_E_*^2^ is the variance of *E*. Since the aim of GWAS is to detect any association between SNPs and a given trait, a shift in the phenotypic distributions or variance is of interest. The conventional association test which targets the location shift only is not adequate for this purpose. Instead, tests targeting the location and scale shifts jointly or, more generally, distributional shifts can capture more general genotype effects. Furthermore, Soave et al. (2015) showed through simulations that these tests can achieve a superior power compared with the conventional test in the presence of an unknown genetic interaction.

### Joint location and scale tests and rank-based inverse normal transformation

We provide a review of four joint location and scale tests, likelihood ratio test (LRT) (Cao et al., 2014); double generalized linear model (DGLM) (Rönnegård and Valdar, 2011); joint location-scale (JLS) test (Soave et al., 2015); and joint location-scale score (JLSS) test (Staley et al., 2021), in Appendix A. These differ generally by the estimation of scale parameters, the iterative process to fit the joint models and ability to account for the correlation between location and scale parameters estimates. These joint location and scale tests require normality of the error term. When this assumption is violated, a common remedial measure is to transform the data. Rank-based inverse normal transformation is a popular data transformation technique in genetic epidemiology, mapping a variable into another that is perfectly normal. The idea of this data transformation is to estimate the empirical quantiles of the variable of interest through its fractional ranks and to apply a quantile function of the normal random variable. The INT is defined as follows:

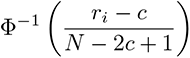

where *N* is the sample size, *r_i_* is the fractional rank of the *i^th^* observation (*i.e.* rank of *i^th^* observation / *N*), *c* ∈ [0, 1] is an offset term and Φ(·) is the quantile function of the standard normal random variable. Since fractional ranks of 0 or 1 would yield −∞ or +∞ after the application of the quantile function, respectively, the role of the linear transformation with the offset *c* is to avoid a value of 0 or 1. A popular offset value is *c* = 3*/*8 (Blom, 1958) but other alternatives exist, such as *c* = 0 (Van der Waerden, 1952), *c* = 1*/*3 (Tukey, 1962) and *c* = 1*/*2 (Bliss et al., 1967). The choice of the offset value is immaterial since these are approximately a linear transformation of each other and close to the expected normal scores (Beasley et al., 2009; Tukey, 1962).

#### Direct INT

Applying the INT directly to the marginal phenotype is referred to as the direct INT (D-INT) (McCaw et al., 2020). This is perhaps the most popular transformation in GWAS among the different forms of INT since the transformed phenotype perfectly follows the standard normal distribution. The steps for analyses with D-INT are highlighted as follows:

1. Apply the INT to *Y* to obtain *Y_T_* .
2. Apply the regression model using *Y_T_* as the response and *G* and ***Z*** as predictors.

Although the idea may appear appealing, the D-INT does not guarantee that the normality assumption is met since this distributional assumption in regression analysis applies to the error term, not to the marginal response variable. For this reason, a second approach, indirect INT, is considered.

#### Indirect INT

The indirect INT (I-INT) consists of applying the INT to the residuals of the phenotype after regressing on the covariates ***Z*** which we denote by *R_Y |_****_Z_***. We review two variants of this type discussed in McCaw et al. (2020). The first I-INT which we refer to as single-adjusted INT (IINT1) is using *R_Y |_****_Z_*** as the response in the regression model with *G* and ***Z*** as predictors. The following steps implement the I-INT1:

1. Regress *Y* on ***Z*** and obtain the residuals *R_Y |_****_Z_***.
2. Apply the INT to *R_Y |_****_Z_*** to obtain *Y_T_* .
3. Apply the regression model using *Y_T_* as the response and *G* and ***Z*** as predictors.

We note that step 1 is to orthogonalize the phenotype *Y* with respect to the space spanned by the columns of ***Z*** and, therefore, ***Z*** should be omitted from the regression model in step 3. However, due to the non-linear change by INT in step 2, the residuals *R_Y |_****_Z_*** are not perfectly orthogonal to the space spanned by the columns of ***Z***, which explains the second adjustment for ***Z*** in step 3 (Sofer et al., 2019).

A second type, referred to as double-adjusted INT (I-INT2), consists of orthogonalizing both the phenotype *Y* and genotype *G* with respect to the space spanned by the columns of ***Z***, yielding *R_Y |_****_Z_*** and *G_T_*, respectively. The INT is applied to the residualized phenotype *R_Y |_****_Z_*** yielding *Y_T_*.

Then, a regression model can be applied using *Y_T_* as the response and *G_T_* as the predictor, omitting ***Z***. The steps for analyzing data with I-INT2 are shown as follows:

1. Regress *G* on ***Z*** and obtain the residuals *G_T_* .
2. Regress *Y* on ***Z*** and obtain the residuals *R_Y |_****_Z_***.
3. Apply the INT to *R_Y |_****_Z_*** to obtain *Y_T_* .
4. Apply the regression model using *Y_T_* as the response and *G_T_* as the only predictor (omitting ***Z***).

Similar to the I-INT1, the transformed phenotype *Y_T_* is not perfectly orthogonal to the space spanned by the columns of ***Z*** due to the INT in step 3 and, therefore, ***Z*** may be required in the regression model in step 4. However, our simulation studies in Appendix B show that the I-INT2 without this second adjustment has good control of type I error overall.

We note that the extent of the INT’s impact on type I error and power is not well understood. In particular, the INT is a non-linear transformation that changes the phenotypic scale, and, as a consequence, the performance of scale tests would be expected to be affected by INT. We conducted extensive simulation studies to investigate the effect of INT on the joint location and scale tests. The results showed that applying I-INT2 leads to a well-controlled type I error and can improve the power under non-normality. The details are provided in Appendix B. However, we are unsure how generalizable these results are outside of the specific simulation scenarios investigated. Thus, in the next section, we propose a new robust approach in which the type I error does not depend on a prior distributional assumption and is as powerful as other location and scale tests under normality. We used quantile regression as the test framework, hence, we refer to this test as the quantile regression-based joint location-scale (qJLS) test.

We note that there exists non-parametric and semi-parametric tests which are robust, with the type I error unaffected by non-normal error distributions. However, when the error distributions are moderately close to a normal distribution, these show a deficit in power compared with the joint location and scale tests. In addition, these methods are computationally inefficient and cannot adjust for continuous covariates, which may be confounders such as principal components for genetic ancestry.

### Quantile regression-based joint location-scale test (qJLS)

Consider the following linear quantile regression model:

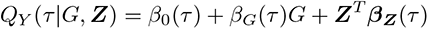

where *Q_Y_* (*τ|G,* ***Z***) is the *τ* ^th^ quantile of the phenotype *Y* for a given SNP *G* and *p*-dimensional covariates ***Z***. The parameters *β*_0_(*τ*), *β_G_*(*τ*) and ***β_Z_***(*τ*) are the quantile-specific intercept, effect of *G* and effects of ***Z***, respectively. Since the unknown genetic interaction distorts the conditional phenotypic distribution across the genotypic groups, we are interested in detecting any quantile-specific effect of *G* across the quantiles. In other words, the hypothesis of interest is the following:

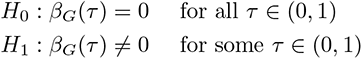

Under some conditions, Koenker and Bassett (1978) provided the asymptotic distribution of *L*-variate estimators 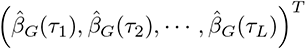 under the null hypothesis, as follows:

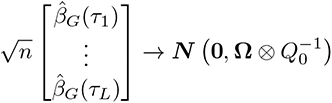

where **Ω** is a *L × L* matrix with the following elements:

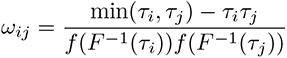

with *F*, the distribution function of *ε* with associated density *f*, and

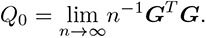

Based on this result, a Wald-type test can be constructed. However, the asymptotic variance-covariance matrix is a function of the density function of the error *ε*. Due to the added uncertainty in estimating this density, the Wald-type test can show an inflated type I error, as seen in our preliminary simulation study (Supporting Information Table S8). Alternatively, robust alternatives such as kernel-based estimators can be considered. In our preliminary simulation studies, although these robust estimators showed a reduction in the type I error, an inflation in type I error occurred at quantiles where the data are sparse or when the sample size is relatively small, consistent with the findings by Song et al. (2017). Estimation based on resampling is another alternative but is computationally inefficient for GWAS. A second challenge with this approach is that one cannot consider all possible *τ* ’s and needs to select quantiles to test. Our preliminary simulation studies revealed that the choice of quantiles affects the power. As a trivial example, when the distributional shift occurs at specific quantiles, the test showed no power if these quantiles were not selected in the test. The joint location and scale test by Koenker and Xiao (2002) alleviates this problem of quantile selection, however, it requires estimation of the error density and the asymptotic null distribution does not have a closed form.

For these reasons, we implement the rankscore test for quantile regression (Gutenbrunner and Jurecková, 1992; Gutenbrunner et al., 1993), which is free of the error density and is computationally efficient. Briefly, suppose the null hypothesis holds (*i.e. β_G_*(*τ*) = 0 for all *τ*) and the true regression model is the following:

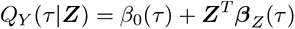

The regression rankscore process for the *i^th^* observation is defined as follows:

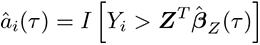

where *I*(·) is the usual indicator function and ***β̂****_Z_* (*τ*) are the regression parameters estimated under the null hypothesis. In other words, after fitting the model under the null hypothesis, if the *i^th^* residual lies above the fitted hyperplane, *â_i_*(*τ*) takes a value of 1 and 0 otherwise. To obtain the regression rankscore *b̂_i_*, *â_i_*(*τ*) are combined over *τ* ∈ (0, 1) with some weighting function *ϕ*(*τ*) called the score function, as follows:

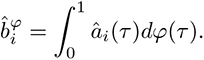

The score function *ϕ*(*τ*) plays a central role because the efficiency of the rankscore test is dictated by its choice. Choosing an optimal score function requires the knowledge of the error distribution and the nature of the distributional shift under the alternative hypothesis. Koenker (2010) considered some useful distributional shifts and provided a list of optimal score functions for a selection of common error distributions. For example, when the error term follows a normal distribution with a constant location shift over all quantiles, the optimal score function is:

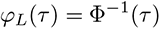

where Φ is the Gauss error function. Another example of interest is when the error follows a normal distribution with a constant scale shift over all quantiles. The corresponding optimal score function for this alternative is:

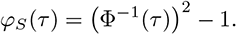

From this, the following test statistic can be computed:

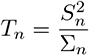

where 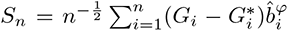 and 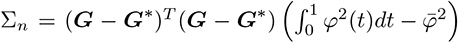. Here, *G^∗^* is the orthogonal projection of G onto the space spanned by the columns of the covariates Z and 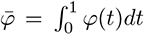. Under the null hypothesis and conditions defined in Appendix C, *T*_n_ converges in distribution to the *χ*^2^(1) distribution. Essentially, the score *S_n_* measures the linear association between (*G−G^∗^*), the covariate-adjusted genotype, and *b̂^ϕ^*, the rankscores under the null hypothesis that are weighted by the selected score function. When no such association exists, we expect *S_n_* to be 0, since the rank of the phenotype is not affected by the genotype.

Furthermore, when considering the alternative hypothesis, one can be specific about the nature of the distributional shift. For example, when a location shift occurs over all quantiles, the corresponding alternative is *β_G_*(*τ*) = *β_G_* for all *τ* . Under a local alternative of the type 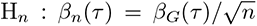, Gutenbrunner et al. (1993) showed that *T_n_* converges in distribution to a non-central *χ*^2^ distribution with 1 degree of freedom and the following non-centrality parameter:

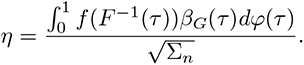

From this result, the test optimality depends on the score function, error distribution and local alternative that is formulated for specific shifts in the distribution. When the error distribution and the nature of the distributional shift is known, the optimal score function can be derived by maximizing the non-centrality parameter (Koenker, 2010). However, the error distribution and the type of distributional shift are generally unknown *a priori*. In the case of GWAS with unknown interactions, we chose to maximize the efficiency for a location and scale shift under normality, and combine both rankscore tests using their joint asymptotic distribution. We refer to this test as qJLS. The score vector of the qJLS test is:

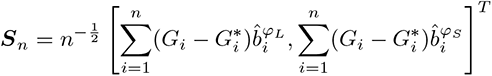

where *ϕ_L_* and *ϕ_S_*, as defined above, are the optimal score functions for the location shift and scale shift, respectively, under a normal error. The following is the test statistic for the qJLS test:

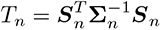

where

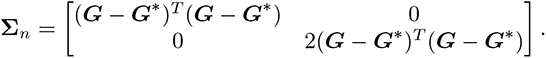

Under the conditions defined in Appendix C, the test statistic *T_n_* converges in distribution to *χ*^2^(2) under the null hypothesis. The proof is provided in Appendix C.

We note that the asymptotic distribution of the test statistic under the null hypothesis does not contain any parameters from the error distribution. Therefore, the type I error is independent of the error distribution. On the other hand, the score functions were chosen based on test optimality under a normal distribution. In other words, under a non-normal error, the type I error of the qJLS test is still well-controlled, but the power of the test is affected due to our choice of the score functions. In comparison, the other joint location and scale tests reviewed in Appendix A assume normality but show an inflated type I error when the error distribution departs from normality (Appendix B). Although I-INT2 showed control of the type I error and/or improvements in statistical power, its statistical properties have yet to be defined formerly since the simulation studies only included a limited number of scenarios. In contrast, qJLS guarantees a well-controlled type I error regardless of the error distribution, a main advantage over the other joint location and scale tests. The qJLS software package is publicly available on Strug Lab, along with the simulation code (https://github.com/strug-hub/qJLS).

## Verification and comparison

We note that most simulation studies in the literature used linear heteroscedastic models as a data generating mechanism, which can only generate a location and/or scale shift. However, such a setting is limited since an unknown genetic interaction may modify the entire shape of the conditional distribution of the phenotype, beyond the location or scale. Here, we used the genetic interaction model (Model 1) as described in Paŕe et al. (2010). We extended the simulation settings by Aschard et al. (2013) to include non-normal errors, quantile-specific interaction effects and unknown background heterogeneity. Lastly, we used linear location and scale models to generate a pure location or scale shift without any genetic interaction, ideal for conventional location testing and scale testing, and to evaluate the loss of power by using joint location and scale tests compared with these conventional tests.

### Power

#### Single unknown interaction

The following is the data generating model:

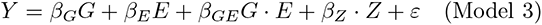

The unknown exposure *E* was a Bernoulli random variable with a probability of success *p_E_* = 0.3. We generated the number of minor alleles *G* with a minor allele frequency of *p* = 0.3 and the covariate *Z* from a standard normal distribution such that these variables were weakly correlated, based on the approach in Cao et al. (2014). Briefly, we first sampled from a bivariate normal distribution 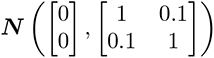. One of these two variables were discretized based on the minor allele frequency *p* = 0.3, yielding *G*.

For each simulation study, we sampled the error term *ε* from (i) standard normal *N* (0, 1); (ii) *χ*^2^(3); and (iii) empirical error distribution from the data application in Section 4. We then standardized the error term by subtracting its expected value and dividing by its standard deviation. The selected density functions of the error are displayed in Figure 2.

**Figure 2:**
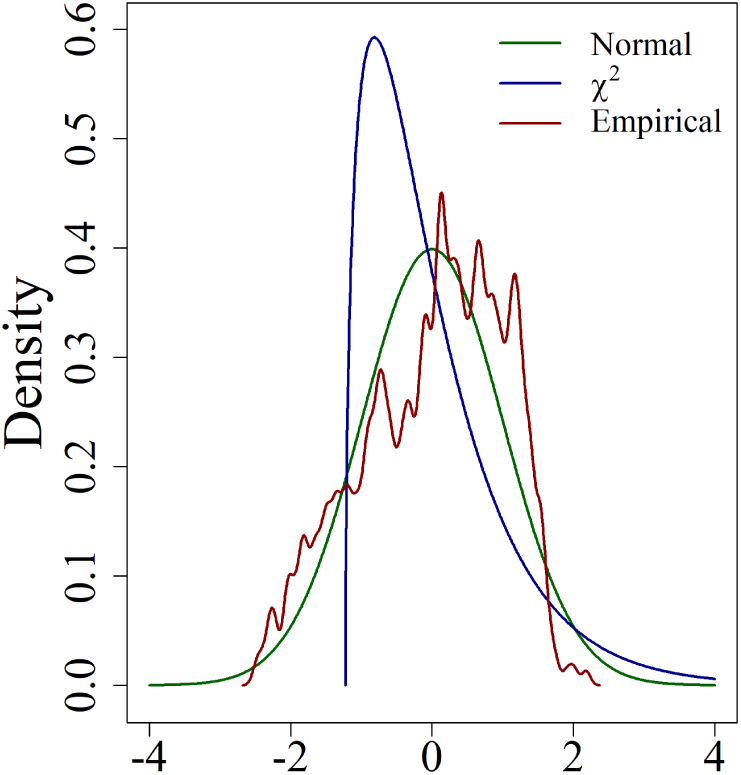
Density functions of the error term *ε* for the simulation models: the selected random variables included standardized normal (in green), *χ*^2^(3) (in blue) and empirical distribution of the residualized top SNP in Section 4 (in red).

By fixing the main effects, we varied the interaction effect *β_GE_* from -1 to 1 by an increment of 0.1 and estimated the power at each increment at the genome-wide significance level of 5 *×* 10*^−^*^8^ based on 100, 000 replicates. We fixed the main effects at *β_G_* = 0.01; *β_E_* = sign(*β_GE_*) *×* 0.3; and *β_Z_* = 0.3.

#### Single unknown quantile-specific interaction

We modified Model 3 to induce an unknown genetic interaction that affects the upper and lower quantiles of the error term as follows:

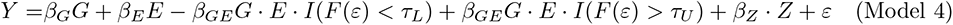

where *F* (*·*) is the distribution function of the error term *ε*, *I*(*·*) is the usual indicator function, and *τ_U_* and *τ_L_* are some upper and lower quantiles, respectively. Essentially, in this setting, the interaction only affects the lower quantile below *τ_L_* with a negative effect *−β_GE_*, and upper quantiles above *τ_U_* with a positive effect *β_GE_*. We fixed *τ_U_* = 0.8 and *τ_L_* = 0.3. The simulation settings were kept the same as in the scenario with a single unknown variable above except that we varied the interaction effect *β_GE_* from 0 to 1 by an increment of 0.1.

#### Two unknown interactions

The following model includes two binary variables *E*1 and *E*2 interacting with *G*:

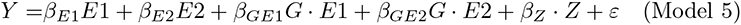

where we sampled *E*1 and *E*2 independently from a Bernoulli distribution with an equal probability of success *p_E_*_1_ = *p_E_*_2_ = 0.3. We considered two scenarios in which (i) *β_GE_*_1_ = 0.2; and (ii) *β_GE_*_1_ = *−β_GE_*_2_. For the former scenario, we varied *β_GE_*_2_ from -1 to 1 by an increment of 0.1 and, for the latter, from 0 to 1 by an increment of 0.1. The main effects were fixed at *β_E_*_1_ = 0.3. The remaining simulation parameters were kept the same as in the scenario with a single unknown interaction.

### Type I error

#### Null hypothesis

We generated data under the following model by applying *β_G_* = *β_GE_* = 0 in Model 3:

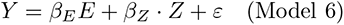

We kept the same setting as in the scenario with a single unknown interaction except that *G* was no longer correlated with *Z* by sampling independently. Since *Z* has an effect on *Y* whereas *G* does not, we removed the correlation between *Z* and *G* to maintain a coherent model. Additionally, for each simulation study, we varied the minor allele frequency (*p* 0.01, 0.05, 0.1, 0.3) and the sample size (*N* 200, 500, 1000, 2000) to study the impact of the minor allele frequency and sample size on the type I error for the tests considered. The type I error was estimated based on 1,000,000 replicates.

#### Null hypothesis with unknown background heterogeneity

Corty and Valdar (2018) considered some unknown source of heterogeneity other than the genetic component, which they termed a “background heterogeneity”, as opposed to the “foreground heterogeneity” that is of researchers’ interests. This nuisance, independent from *G*, may affect the conditional distribution of the phenotype including its location and scale but, since it is unknown, it is left unspecified in the statistical model. For example, forced expiratory volume in 1 second (FEV1) in Canadians with CF is heterogeneous across the age (Kim et al., 2018), hence, age can be a source of background heterogeneity if left unspecified. We study the effect of the background heterogeneity on the type I error with the following model:

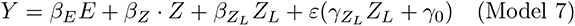

where *Z_L_* is the unknown source of background heterogeneity that can affect the location and/or scale of the phenotype *Y* if *β_ZL_* and *γ_ZL_* are non-zero. We fixed *β_ZL_* = 0.3, *γ_ZL_* = 0.5 and *γ*_0_ = 1, inducing location and scale shifts simultaneously. When estimating the type I error, we omitted *Z_L_* in the model as it is an unknown source. We note that, in this setting, the estimated parameters for the conditional mean and the parameters for the conditional variance are dependent due to their mutual dependence on *Z_L_*.

### Comparison with conventional association tests

In the presence of unknown genetic interactions, joint location and scale tests may have superior power compared with the conventional location or scale association tests. Here, we study the loss of power compared with the conventional association tests when there is no such interaction. We simulated data under a pure location shift as follows:

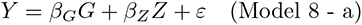

We varied the main effect *β_G_* from -0.5 to 0.5 by an increment of 0.1. The remaining simulation parameters are kept the same as in the scenario with a single unknown interaction. We compared the power of the joint location and scale tests with the classical linear regression which is optimal for this scenario when the error follows a Gaussian distribution.

Similarly, we generated a pure scale shift as follows:

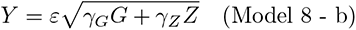

where we applied a standard normal quantile function to *Z* so that *Z ∼* Unif(0, 1). We varied the scale effect *γ_G_* from 0 to 0.5 by an increment of 0.1, keeping other simulation parameters the same as in the scenario with a single unknown interaction. We used the generalized Levene’s test (Soave and Sun, 2017) as the standard and compared the power with the joint location and scale tests.

### Comparison of methods

Using simulation, we compared the performance of the joint location and scale tests (reviewed in Appendix A) with qJLS test. These include the double generalized linear model (DGLM), joint location-scale test (JLS) and joint location-scale score test (JLS-Score). We note that the DGLM, JLS and JLS-Score test require normality of the error term. Since the unknown interaction alters the shape of the distributions, the resulting distribution of the residuals will show a departure from normality based on the normal QQ-plot or through statistical tests, even if the error term in Model 3 follows a normal distribution. On this basis, we applied the I-INT2 as a remedial measure for all error distributions, including the normal distribution since our simulation studies in Appendix B showed that the I-INT2 removed the type I error inflation with skewed or heavy-tailed error distributions and the other variants, D-INT and single-adjusted indirect INT (I-INT1), still resulted in an inflated type I error or in numerical instability in some scenarios. Therefore, we implemented the I-INT2 as the transformation function for the DGLM and JLS tests. Our preliminary simulation results showed that the JLS-Score test remained robust in all simulation scenarios that we considered (Table 4 in Appendix B). However, the power was considerably reduced without any data transformation under the alternative hypothesis in scenarios where the error term departed severely from normality. On this basis, we applied an INT for the JLS-Score test but also considered the case without any data transformation. We chose the I-INT2, solely for comparison purposes. We note that the power of the JLS-Score test varied with the type of INT, error distribution and type of distributional shift in our simulation studies in Appendix B and no single INT led to the highest power. We did not apply any transformation function for qJLS since the asymptotic distribution of its test statistic under the null hypothesis is independent of the error distribution under the null hypothesis and, therefore, the type I error is well-controlled regardless of the error distribution. For the simulation study in Section 3.3 that is based on pure location or scale shift without any interaction, we did not apply any transformation when the error was normal since the normality actually holds in these scenarios. When the error was non-normal, we applied the direct INT (D-INT) to the conventional location or scale tests, since this is commonly used in GWAS, and I-INT2 for the joint location and scale tests except the qJLS test.

### Results

#### Type I error

Table 1 displays the empirical type I error, varying nominal levels and error distributions under Model 6 with minor allele frequency of 0.3 and sample size *N* = 2, 000.

**Table 1:** Empirical type I error under Model 3 varying the error distribution. . The sample size and minor allele frequency (MAF) were fixed at 2,000 and 0.3, respectively. Four tests including double generalized linear model (DGLM), joint location-scale (JLS) test, joint location-scale score (JLS-Score) test and quantile regression based joint location-scale (qJLS) test were used. For the first three tests, we applied no transformation or indirect double-adjusted rank-based inverse normal transformation (I-INT2) to the data. The type I error was estimated based on 1,000,000 replicates.

| Error Distribution | Nominal p-value | No transformation |  |  |  | I-INT2 |  |  |
| --- | --- | --- | --- | --- | --- | --- | --- | --- |
|  |  | DGLM | JLS | JLS-Score | qJLS | DGLM | JLS | JLS-Score |
| Normal | $5 \times 10^{-2}$ | 0.050225 | 0.049610 | 0.049605 | 0.049219 | 0.048864 | 0.049365 | 0.049581 |
| | $5 \times 10^{-3}$ | 0.005031 | 0.004939 | 0.004921 | 0.004848 | 0.004747 | 0.004880 | 0.004916 |
| | $5 \times 10^{-4}$ | 0.000506 | 0.000477 | 0.000480 | 0.000478 | 0.000461 | 0.000472 | 0.000480 |
| | $5 \times 10^{-5}$ | 0.000063 | 0.000061 | 0.000056 | 0.000051 | 0.000053 | 0.000054 | 0.000056 |
| $\chi^2$ | $5 \times 10^{-2}$ | 0.204755 | 0.079244 | 0.049834 | 0.049380 | 0.048960 | 0.049550 | 0.049751 |
| | $5 \times 10^{-3}$ | 0.084472 | 0.018055 | 0.004860 | 0.004745 | 0.004667 | 0.004844 | 0.004797 |
| | $5 \times 10^{-4}$ | 0.037523 | 0.004398 | 0.000473 | 0.000504 | 0.000482 | 0.000483 | 0.000491 |
| | $5 \times 10^{-5}$ | 0.017276 | 0.001057 | 0.000048 | 0.000055 | 0.000048 | 0.000052 | 0.000055 |
| Empirical Data | $5 \times 10^{-2}$ | 0.034136 | 0.058374 | 0.050120 | 0.049429 | 0.048979 | 0.049795 | 0.049976 |
| | $5 \times 10^{-3}$ | 0.003579 | 0.008962 | 0.005043 | 0.004904 | 0.004812 | 0.004933 | 0.004993 |
| | $5 \times 10^{-4}$ | 0.000412 | 0.001525 | 0.000507 | 0.000494 | 0.000472 | 0.000496 | 0.000506 |
| | $5 \times 10^{-5}$ | 0.000051 | 0.000263 | 0.000050 | 0.000047 | 0.000040 | 0.000050 | 0.000044 |

Under the standard normal distribution, the type I error estimates for all joint tests without any data transformation are close to each nominal level, as expected. However, the estimated type I error rate for the DGLM was slightly above all nominal levels. Under the skewed *χ*^2^ distribution, the inflation was evident for the DGLM with a large empirical type I error, as well as the JLS test that showed a relatively moderate inflation. For both methods, the inflation was corrected after applying the I-INT2. The histograms of p-values with I-INT2 appeared to be uniform, further supporting the correction by the data transformation (Supporting Information Figure S4). For the JLS-Score test, the data transformation was unnecessary since the empirical type I error was below the nominal levels without the I-INT2. Under the empirical data distribution, the empirical type I error for DGLM was well below the nominal levels. The histogram of p-values shows a skewness to the left, indicating that the method is overly conservative for the given distribution (Supporting Information Figure S5). After applying I-INT2, the histogram of p-values appeared uniform. A moderate inflation was noted for the JLS test but was corrected after applying the I-INT2. The JLS-Score test remained robust in this scenario with empirical type I error below the nominal levels. In all scenarios, qJLS did not show any evidence of inflation, as expected.

When the sample size was reduced, estimated type I error increased for the DGLM under the standard normal distribution whereas other methods were unaffected (Supporting Information Table S1). This behaviour for DGLM is consistent with the previous simulation studies by Soave and Sun (2017). The empirical type I error for the DGLM was slightly above the nominal levels for all sample sizes. At the nominal level of 0.05, the estimated type I error was above this level for all sample sizes except *N* = 2, 000. Furthermore, when *N* = 200, approximately 0.011% of the replicates did not reach the numerical convergence. After applying the I-INT2, the empirical type I error for the DGLM fell below the nominal levels. The transformation resolved the optimization problem with 100% of the replicates attaining the convergence. Under *χ*^2^, the DGLM showed a large empirical type I error and increased failures in numerical convergence, close to 0.37% when *N* = 200, which were resolved by applying the I-INT2 (Supporting Information Table S2). A moderate inflation was noted for the JLS test, which was corrected after applying the I-INT2. The results for the empirical data distribution followed the pattern observed when *N* = 2, 000 (Supporting Information Table S3). The empirical type I error for the JLS-Score and qJLS tests did not show any sign of inflation with the change in the sample size.

The reduction of minor allele frequency affected the empirical type I error for all methods except the JLS test under the standard normal distribution (Supporting Information Table S4). For the DGLM, type I error estimates were slightly above the nominal levels when MAF = 0.01. For the JLS-Score and qJLS tests, when the minor allele frequency decreased, the inflation increased at the tail of the null distribution, although at the nominal level of 0.05 no inflation was observed. In all cases, applying the I-INT2 did not correct the inflation. A similar trend is observed under the *χ*^2^ and empirical data distribution (Supporting Information Table S5 and Table S6, respectively). When the minor allele frequency decreased, a slight inflation was observed for the DGLM with I-INT2 whereas this inflation occurred at the tail of the null distribution for the JLS-Score test with I-INT2 and the qJLS test. There was no evidence of inflation for the JLS test with I-INT2 in any of the scenarios.

Table S7 (Supplementary Information) displays the effect of an unknown background heterogeneity affecting simultaneously the location and scale of the phenotypic distribution on type I error, which results in a dependence between the estimators for the location parameters and the ones for the scale parameters. Since the DGLM and JLS test rely on the independence of estimators for the location and scale parameters, both methods were affected by the background heterogeneity. The DGLM exhibited a large inflation for all error distributions that included the standard normal distribution whereas the inflation was moderate for the JLS test. For both methods, the inflation was corrected after applying the I-INT2. The JLS-Score and qJLS tests remained robust to the unknown background heterogeneity.

#### Power

Figure 3 shows the estimated power under a single unknown interaction model (Model 3) as a function of the interaction effect *β_GE_* based on minor allele frequency of 0.3, sample size *N* = 2, 000 and 100,000 replicates at each value of *β_GE_*. Under the standard normal distribution, the power of the qJLS test was slightly greater than the other methods that required the I-INT2. Among these three methods that required the I-INT2, no difference in power was discernible. Under the *χ*^2^ distribution, the qJLS test showed the greatest power when the interaction effect was negative, closely followed by the DGLM. When the interaction effect was positive, JLS and JLS-Score tests showed greater power than qJLS and DGLM. Under the empirical data distribution, the qJLS test was the most powerful for negative and positive values of *β_GE_*.

**Figure 3:**
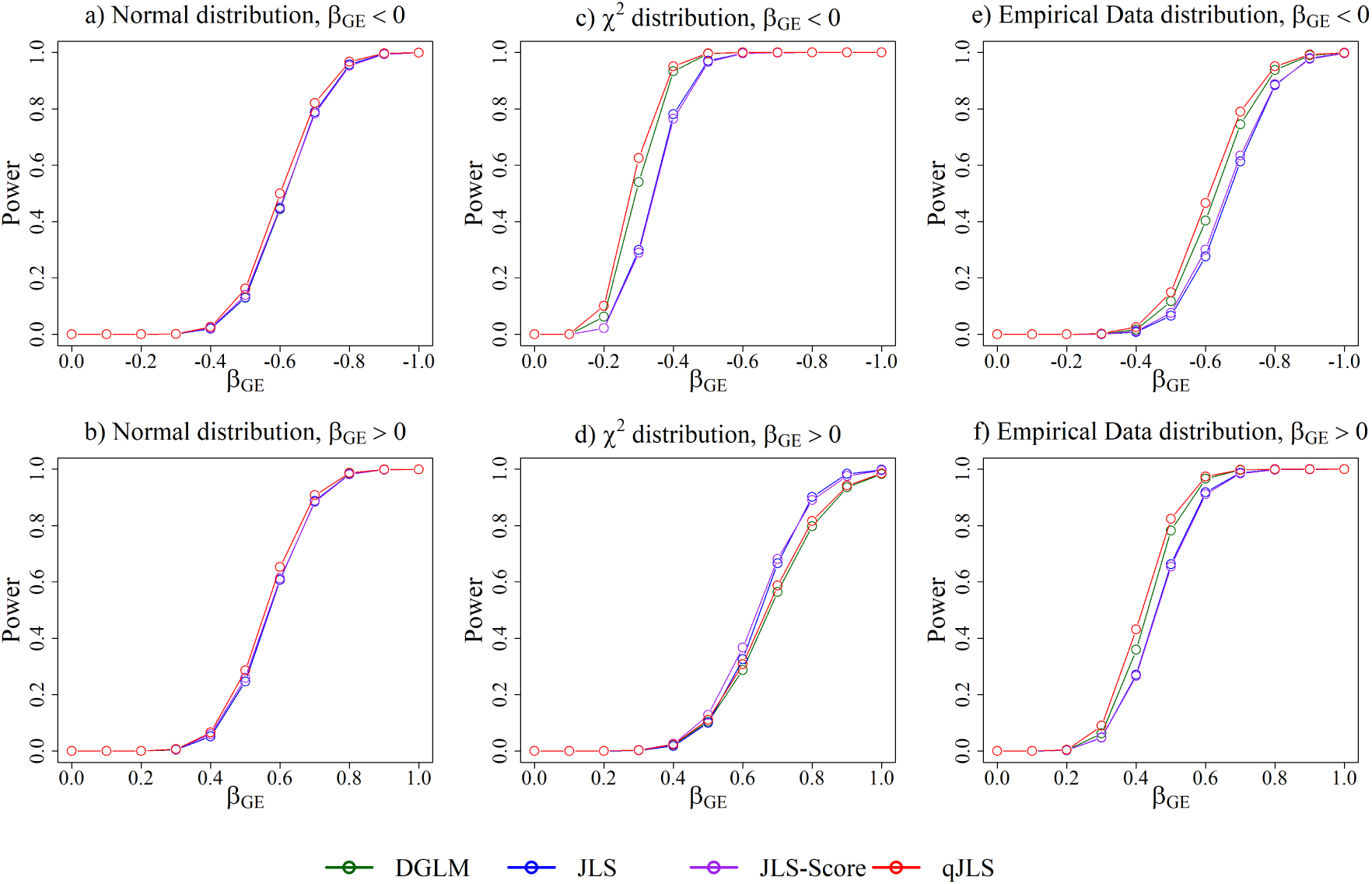
Empirical power under an unknown genetic interaction model with a binary single variable (Model 3) by error distribution: the strength of the interaction effect *β_GE_* was varied. The top and bottom rows display negative and positive interaction effects, respectively. The error distributions, displayed in each column, included the standardized normal, *χ*^2^(3) and empirical distribution of the residualized top SNP in Application section. The sample size and minor allele frequency (MAF) were fixed at 2,000 and 0.3, respectively. Four tests including double generalized linear model (DGLM in green), joint location-scale (JLS in blue) test, joint location-scale score (JLS-Score in purple) test and quantile regression based joint location-scale (qJLS in red) test were used. For the first three tests, we applied the indirect double-adjusted rank-based inverse normal transformation (I-INT2) to the data for all error distributions. The power was estimated based on 100,000 replicates with the nominal level 5 *×* 10*^−^*^8^.

Figure 4 shows the estimated power under the quantile-specific unknown interaction model (Model 4). The qJLS test showed the greatest power for all error distributions. Interestingly, the power of the JLS and JLS-Score tests was approximately 0 for all error distributions considered, showing that the JLS and JLS-Score tests were not able to detect these distributional changes at the tail.

**Figure 4:**
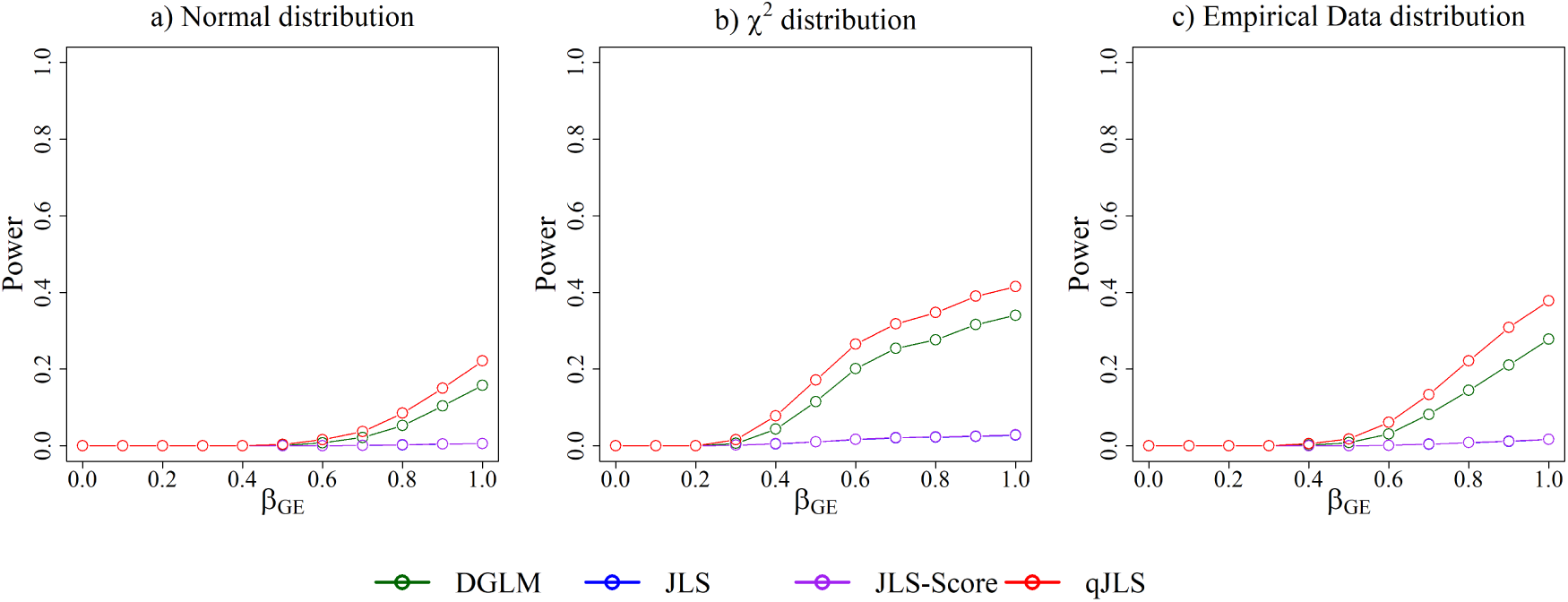
Empirical power under a quantile-specific unknown genetic interaction model with a binary single variable (Model 4) by error distribution: the unknown interaction effect *β_GE_* was varied, affecting the upper quantile of the error term above 0.8 positively with *β_GE_ >* 0 and the lower quantile below 0.3 negatively with *β_GE_*. The error distributions, displayed in each column, included the standardized normal, *χ*^2^(3) and empirical distribution of the residualized top SNP in Application section. The sample size and minor allele frequency (MAF) were fixed at 2,000 and 0.3, respectively. Four tests including double generalized linear model (DGLM in green), joint location-scale (JLS in blue) test, joint location-scale score (JLS-Score in purple) test and quantile regression based joint location-scale (qJLS in red) test were used. For the first three tests, we applied the indirect double-adjusted rank-based inverse normal transformation (I-INT2) to the data for all error distributions. The power was estimated based on 100,000 replicates with the nominal level 5 × 10*^−^*^8^.

In Figure 5, the power was estimated for two interacting variables among which one was fixed with a weak effect of 0.2 and the interaction effect of the other was varied, based on Model 5. For the normal error, the estimated power was the greatest for the qJLS test. A similar pattern emerged for the *χ*^2^ and empirical data distributions.

**Figure 5:**
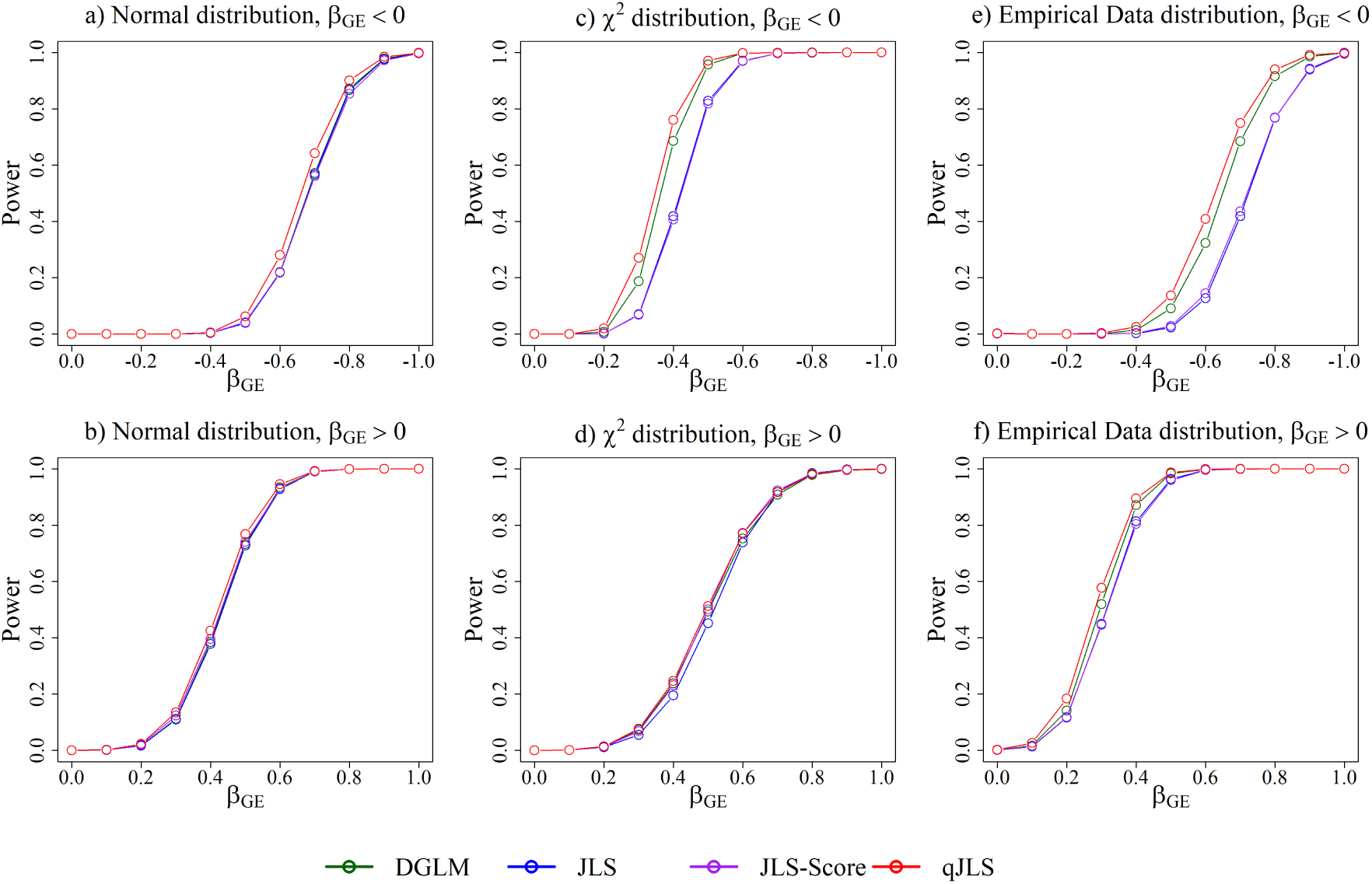
Empirical power under an unknown genetic interaction model with two independent binary variables, *E*1 with fixed effect, and *E*2 with varying effects, (Model 5) by error distribution: after fixing the interaction effect of *E*1 at *β_GE_*_1_ = 0.2, we varied the interaction effect of *E*2, *β_GE_*_2_. The top and bottom rows display negative and positive interaction effects of *E*2, respectively. The error distributions, displayed in each column, included the standardized normal, *χ*^2^(3) and empirical distribution of the residualized top SNP in Application section. The sample size and minor allele frequency (MAF) were fixed at 2,000 and 0.3, respectively. Four tests including double generalized linear model (DGLM in green), joint location-scale (JLS in blue) test, joint location-scale score (JLS-Score in purple) test and quantile regression based joint location-scale (qJLS in red) test were used. For the first three tests, we applied the indirect double-adjusted rank-based inverse normal transformation (I-INT2) to the data for all error distributions. The power was estimated based on 100,000 replicates with the nominal level 5 *×* 10*^−^*^8^.

Figure 6 displays the estimated power for two interacting variables with opposite interaction effects, based on Model 5. The qJLS test showed the greatest power for all error distributions. For the *χ*^2^ and empirical data distribution, the difference in the power between the qJLS test and the other three tests was more pronounced.

**Figure 6:**
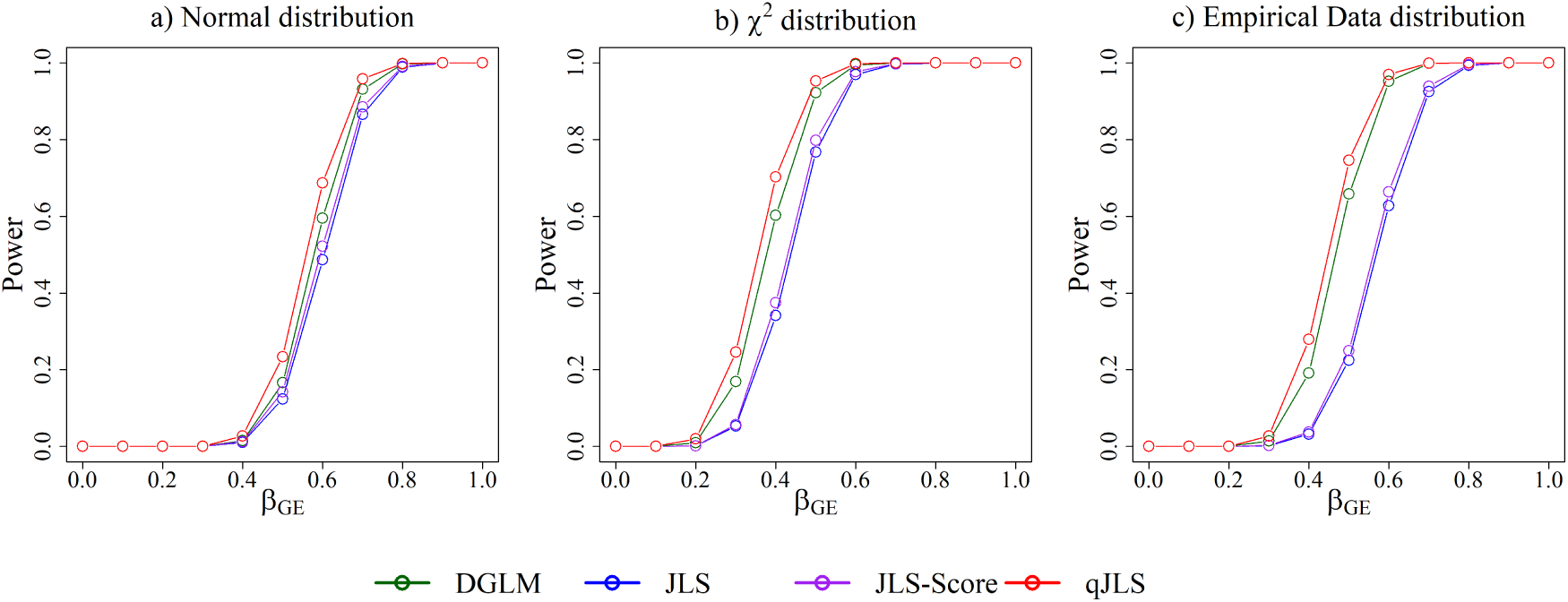
Empirical power under an unknown genetic interaction model with two independent binary variables, *E*1 and *E*2 with opposite effects, (Model 5) by error distribution: the interactions effects of *E*1 and *E*2 were opposite *i.e. β_GE_*_1_ = *βGE*2. The error distributions, displayed in each column, included the standardized normal, *χ*^2^(3) and empirical distribution of the residualized top SNP in Application section. The sample size and minor allele frequency (MAF) were fixed at 2,000 and 0.3, respectively. Four tests including double generalized linear model (DGLM in green), joint location-scale (JLS in blue) test, joint location-scale score (JLS-Score in purple) test and quantile regression based joint location-scale (qJLS in red) test were used. For the first three tests, we applied the indirect double-adjusted rank-based inverse normal transformation (I-INT2) to the data for all error distributions. The power was estimated based on 100,000 replicates with the nominal level 5 *×* 10*^−^*^8^.

#### Comparison with conventional association test

Figure S1 (Supporting Information) shows the comparison of the power between the classical linear regression and the joint location and scale tests under a pure location shift model without any interaction (Model 8 - a), varying the main genetic effect and error distributions. In the scenario with the standard normal distribution, the classical regression is known to yield the best linear unbiased estimator. This theory is well demonstrated in Figure S1 (Supporting Information) in which linear regression was the most powerful, although the power reduction for the joint location and scale tests was moderate. The average difference over all effect sizes considered was 3% with the maximum difference of 11%. Further simulation studies showed that the qJLS test required approximately a 12% increase in sample size to achieve similar power under these conditions. When considering *χ*^2^, all joint location and scale tests showed a large improvement in the power compared with the classical regression, clearly demonstrating the benefits in this scenario.

Similarly, the generalized Levene’s test for the variance heterogeneity was compared with the joint location and scale tests under a pure scale shift model (Model 8 - b), by varying the main variance effect of *G* and error distributions (Supporting Information, Figure S2). Surprisingly, DGLM and qJLS tests showed the greatest power with a large improvement over the generalized Levene’s test under the standard normal distribution. The difference in power between DGLM and qJLS was minimal. JLS and JLS-Score tests had a moderately inferior power compared with the generalized Levene’s test. For the remaining error distributions, qJLS showed a large positive difference compared with the remaining tests. No difference in the power for the tests except qJLS can be discerned from Figure S2 (Supporting Information).

## Application

### Method

#### Study sample

The Canadian CF Gene Modifier Study (CGMS) was designed to recruit a representative sample of the Canadian CF population with the objective to identify genes that modify CF disease severity across the affected organs and notably in the lungs (Taylor et al., 2006). Here we investigate the use of joint location scale tests to identify loci that modify lung disease severity. The study was approved by the Research Ethics Board of the Hospital for Sick Children and written informed consent was obtained from each participant.

Clinical data were obtained via chart review and through the Canadian CF Registry, which captures longitudinal follow-up of the Canadian CF population.

#### Pulmonary phenotype

We used the longitudinal spirometry data in the last three years preceding the enrollment date for each participant enrolled in the CGMS whose genome-wide genotype data were available. If the phenotype data were missing prior to the enrollment dates, we used the closest 3-year range of data after the enrollment date. The analysis was limited to individuals with at least 2 or more visits after the age of 6 years for reliable spirometric measurements. Additionally, we only included individuals with two severe CF transmembrane conductance regulator (*CFTR*) mutations associated with pancreatic insufficiency (PI) (Corey et al., 1997). Measurements after transplantation, diagnosis of chronic *B. Cepacia*, or treatment with a CFTR modulator were removed. We computed the CF pulmonary phenotype Saknorm (Taylor et al., 2011), which is the standard normal quantile of forced expiratory volume in 1 second percentiles adjusted for age, sex, height and cohort-specific survival. To generate FEV1-percentiles adjusted for age, sex and height, we used the CF-specific reference equations by Kulich et al. (2005) based on the US CF Foundation (CFF) Registry from 1999 to 2006 for the cohort enrolled in CGMS before 2008 and the Canadian CF-specific reference equations (Kim et al., 2018) based on the Canadian CF Registry data from 2008 to 2014 for those enrolled after 2008. To avoid any extrapolation, we removed individuals with height outside 5cm beyond the height range for the CF-specific reference equations. We then adjusted for the cohort-specific survival probability for each FEV1-percentile and averaged over each individual. We note that we did not apply the standard normal quantile function to this phenotype as in Taylor et al. (2011) since our statistical method qJLS does not require normality. For DGLM, JLS and JLS-Score tests that require normality, we applied I-INT2.

#### Genome-wide genotype data

Genotyping was performed on four Illumina platforms 610Quad, 660W, Omni 2.5 and Omni 5. The detailed quality control (QC) and imputation procedures are described in Gong et al. (2019). SNP position and annotation information were based on Genome Reference Consortium 37 (GRCh37). We included unrelated participants to satisfy the assumption of qJLS, by randomly sampling one individual from each set of related individuals. Additionally, in a principal component analysis (Gogarten et al., 2019), observations inside 6 standard deviations from the centre of African (AFR) or East Asian (EAS) clusters using the 1000 Genomes Project data were excluded from the analysis.

#### Association analysis

The genome-wide association study (GWAS) was performed using the qJLS test. For comparison, we used DGLM, JLS and JLS-Score tests after applying the I-INT2. We used dosage data assuming an additive effect and included the sex, type of reference equations used to compute Saknorm, genotyping platform and 9 principal components as covariates. To account for multiple hypothesis testing, we used the genome-wide significance threshold of 5×10*^−^*^8^ (Dudbridge and Gusnanto, 2008).

## Results

After applying the inclusion-exclusion criteria above, a total of 1,997 participants with CF were included in the analysis. After standard QC and imputation (Gong et al., 2019; Panjwani et al., 2018), 5,533,051 variants were analyzed for association with CF lung disease using Saknorm.

One locus (minimal p-value = 3.12×10*^−^*^8^ at rs9513900 on chr13:102,090,156; MAF=0.31), annotated between *ITGBL1* (chr13:102,105,026-102,373,206) and *NALCN* (chr13:101,706,128-102,068,859) was significantly associated with Saknorm using qJLS (Figure 7).

**Figure 7:**
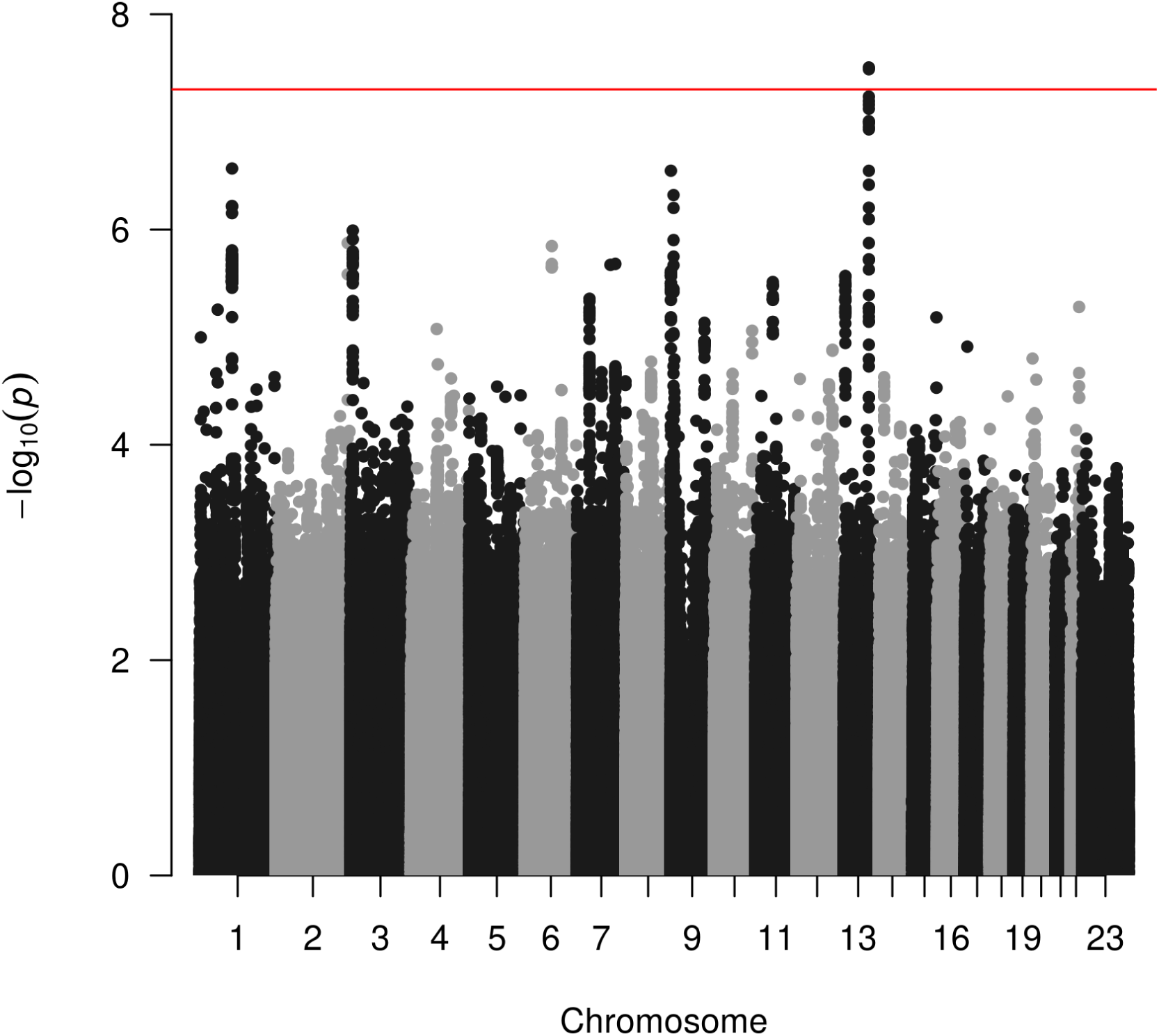
Manhattan plot for the genome-wide association study of Canadian CF lung disease with unrelated sample of 1,997 individuals in CGMS based on qJLS: All variants with MAF *>* 1% are included in the analysis. The solid horizontal line in red corresponds to the genome-wide significance threshold of *p* = 5 *×* 10*^−^*^8^. The top SNP is the variant rs9513900.

Examining the individual location p-value and scale p-value from a linear regression and generalized Levene’s test (without I-INT2), respectively, the association for this SNP was driven by a location shift (*p* = 2.15×10*^−^*^8^) with a positive effect of 0.046 but no significant change in the phenotypic scale detected (*p* = 0.12). This locus was also supported by DGLM, JLS and JLS-Score tests following I-INT2, with this locus showing the smallest p-value genome-wide. However, at the genome-wide significance level, these three other association tests failed to detect any significant loci (Supporting Information, Figure S7). No evidence of inflation in type I error was observed based on the histogram of p-values (Supporting Information, Figure S8) and the QQ-plots (Supporting Information, Figure S9) for all methods.

This genome-wide signficant variant, rs9513900, was not identified in a previous meta-GWAS of CF lung disease (Corvol et al., 2015) that included a subset of the participants analyzed here, with the effect of the variant reported to be 0.033, close to the estimated effect in this study, but did not reach genome-wide significance (p=0.037). Figure S10 (Supporting Information) provides the evidence for this variant across Canadian subsets including data not previously published, all with similar effect size and direction.

Since this variant is intergenic and not in linkage disequilibrium (LD) with any protein coding variants, we assessed whether rs9513900 (or a variant in LD with it) showed evidence of being an expression quantitative trait locus (eQTL) in lung tissue. To determine whether the identified variant rs9513900 shows evidence of an eQTL in lung tissue, we used the data from the genotype tissue expression (GTEx) project (Lonsdale et al., 2013) version 8. Among 12 genes annotated to the region, a significant eQTL for *NALCN* was observed after Bonferroni correction (*p* = 1×10*^−^*^3^). To inspect a possible heterogeneity in the variant’s effects on the phenotype, we applied quantile regression to estimate the quantile-specific effects (Figure 8). Overall, the variant has a positive effect at all percentiles of the conditional phenotype but this effect is differential over *τ*, with the strongest positive effect of 0.074 attained at *τ* = 0.287 that decreases to 0 when approaching the tails. The median effect was 0.054 and was close to the mean effect 0.046 which can be interpreted as the average of the quantile-specific effects over *τ* . Having one variant at this locus increases the median lung phenotype by 5.4% assuming all other predictors in the statistical model are held fixed.

**Figure 8:**
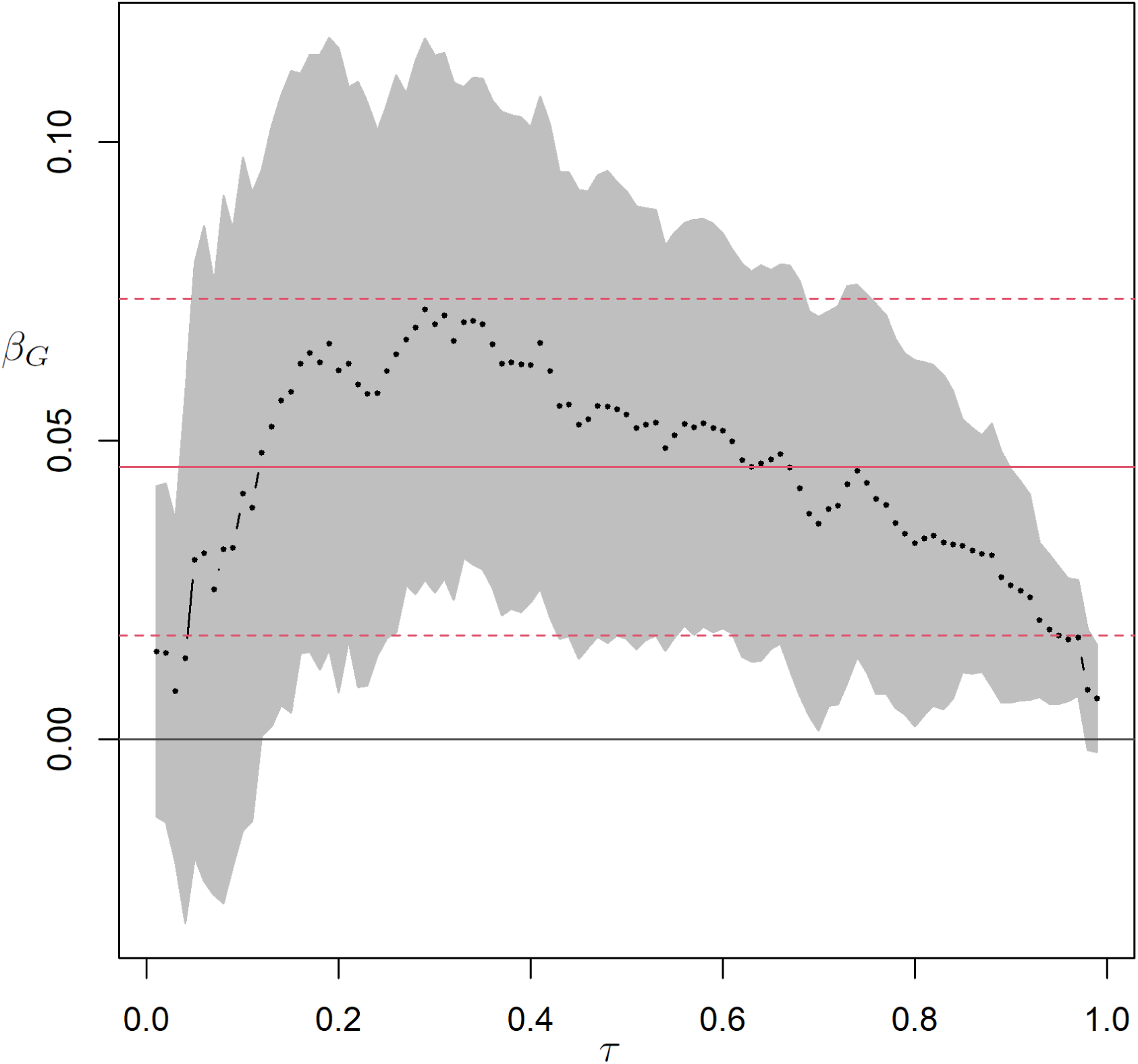
Estimated effects *β*^^^(*τ*) of the variant rs9513900 on the CF pulmonary phenotype at different percentiles. *τ* : quantile regression model was used with dosage data assuming an additive effect. Predictors included the sex, type of CF-specific reference equations used to compute the pulmonary phenotype, genotyping platform and 9 principal components. The shaded gray area represents the simultaneous confidence band using Bonferroni correction. The solid red line is the mean effect, with dashed red line displaying the confidence band.

## Discussion

In this study we proposed a new joint location and scale test, qJLS, based on quantile regression and compared its performance to a selection of joint location and scale tests in the literature. The main benefit of the qJLS test is its robustness to the underlying error distribution while remaining the most powerful under normality. This robustness feature is due to the asymptotic distribution of the test statistic that is completely free of the parameters for the error distribution, under the null hypothesis. In contrast, the joint location and scale tests in the literature are based on a normality assumption which necessitates transformation, such as the rank-based inverse normal transformation, for type I error rate control and improvement in power. Although our simulation studies provided some support for type I error rate control with indirect double-adjusted rank-based inverse normal transformation, this was under specific scenarios and generalization of the results requires a theoretical proof which is currently lacking.

In terms of power, we purposely designed our test to achieve optimality under normality. But, when this assumption fails, the qJLS test was still more powerful than the other methods in most simulation scenarios that were considered. The basis for this performance is mainly attributable to the scale component of the test. We observed that the scale component of the qJLS test was more powerful than that of the other joint location and scale tests while the power for the location component remained similar for all tests considered. This difference was clearly demonstrated when we considered the tail-specific interaction in Figure 4. However, the power of a test is a function of the nature of the distributional shift under the alternative and the error distribution. Since the error distribution differs between phenotypes and unknown genetic interactions may affect the error distribution in different ways, the test optimality may be scenario-specific, which makes it difficult to determine an optimal test *a priori* for real-life data. This is demonstrated in Figure 3 in which qJLS was less powerful than JLS and JLS-Score tests for negative interaction effects under the standardized *χ*^2^(3) distribution. Although we selected rankscore functions that are optimal for location and scale shifts under normality, there are alternative rankscore functions that are optimal for other types of distributional shifts and error distributions (Koenker, 2010). We conducted additional simulation studies based on Model 3, to experiment with alternative rankscore functions defined in Koenker (2010) such as Lehmann, Wilcoxon and trimmed Wilcoxon functions that included either the upper quantiles above the median or lower quantiles below the median, and compared these with the qJLS test (Figure S6 in Supporting Information). The qJLS test performed well overall but, in some scenarios, its power was lower than some alternative rankscore tests. However, the performance of these alternative rankscore tests was extreme, with power close to 0 in some scenarios. Based on these findings, we concluded that qJLS would offer good performance over the range of observable data.

We also explored the benefit of using qJLS over the conventional location or scale test. This benefit was clearly demonstrated in the presence of unknown genetic interactions. Since unknown genetic interactions shift the conditional phenotypic distribution, qJLS harnesses superior power by leveraging both location and scale shifts, rather than each shift individually as in the conventional tests. When we considered a pure location or scale shift without any unknown genetic interaction, qJLS was still more powerful than individual location or scale test except when we considered a pure location shift model with normality. This is the ideal scenario for linear regression, however, the difference in power appeared moderate. Based on simulated data with Model 8 - A, the qJLS test required approximately a 12% increase in sample size to achieve similar power under these optimal conditions for the linear regression. Given the overall benefit, we recommend the use of qJLS over the conventional tests.

The benefit of the qJLS was well-illustrated by the GWAS and its corresponding results. First, we note that, with 5,533,051 variants, the qJLS test demonstrated the fastest computational time. This was expected since this test computes the computation-heavy rankscores only once under the null hypothesis while the remaining iterations for each of the 5,533,051 variants consists of simple multiplications and sums. Second, our qJLS test detected the candidate variant despite exerting a location shift without a significant shift in the phenotypic scale; its p-value (*p* = 3.12×10*^−^*^8^) was only slightly higher than that from the linear regression (*p* = 2.15×10*^−^*^8^). Examination of the identified genome-wide locus suggested the SNP is an eQTL for *NALCN*, a sodium leak channel non-selective protein. NALCN regulates epithelial cell trafficking to distant tissues, including the lung epithelium (Rahrmann et al., 2022). Previous findings (Corvol et al., 2015; Gong et al., 2019; Panjwani et al., 2018; Soave and Sun, 2017) have suggested CF lung disease modifiers impact severity through their role in gene expression variation, providing additional support for the locus, although replication in an independent sample is necessary to establish this gene as a modifier. We also examined percentile-specific effects which were all positive, indicating the variant provides a protective effect at all percentiles of the lung phenotype. But these effects were not uniform across the percentiles and appeared to be quadratic. At both extremes of the phenotype, around 1% and 99%, the effect was nearly zero and achieved its maximum effect at the 29th percentile. This differential effect suggests potential interactions at play, even though no scale shift was detected. A possible explanation for the failure to detect the scale shift is that the current sample size was not sufficient to detect the given magnitude of the scale shift.

Our study provides a review and a comparison of performance across a selection of joint location and scale tests that rely on normality, based on simulation studies. Our findings demonstrate that the DGLM which uses the squared residuals for the scale model was more sensitive to departures from normality than the JLS and JLS-Score that use the robust absolute deviations, without any data transformation. As a clear example, DGLM displayed an inflated type I error even with a logistic distribution which is often indistinguishable from a normal distribution using a QQ-plot or statistical test for normality such as Kolmogorov-Smirnov test. The DGLM is known to suffer from numerical instability with low minor allele frequency (Dumitrascu et al., 2019). In our experience, this instability increased with additional factors such as skewed and heavy-tailed error distributions, low sample size and large numbers of predictors. We showed that applying the indirect double- adjusted rank-based inverse normal transformation fixed the type I error inflation and numerical instability, for the scenarios considered. The JLS test without any data transformation was more robust to distributional misspecifications but showed a moderate inflation in the type I error rate with skewed or heavy-tailed distributions. This inflation was corrected by applying the I-INT2 for the homoscedastic model but, with variance heterogeneity, the JLS test still displayed a slight inflation in type I error, possibly due to the independent estimation of the location and scale parameters. We note that the JLS-Score test which accounts for the correlation between the estimated location parameters and scale parameters remained robust in the simulation scenarios investigated. Although the type I error remained well-controlled, it is the power that suffered when the normality assumption was violated. Applying the I-INT2 resulted in a large improvement in power but the theoretical properties with the data transformation are unknown and remain to be investigated.

We note that a limitation of qJLS is that it is designed for independent observations and cannot accommodate related individuals or repeated observations. A strategy to meet this requirement is to keep one individual per cluster of related individuals, which results in an undesirable reduction in the sample size. Future research will extend qJLS to account for correlation in the data.

## Acknowledgements

We thank the patients, care providers and clinic research assistants, collaborators, and principal investigators involved in CF Centers throughout Canada for their contributions to the CF Canada Patient Registry and Canadian Gene Modifier Study. Funding was provided by Cystic Fibrosis Foundation (STRUG17PO); Cystic Fibrosis Canada (2626); CFIT Program funded by the SickKids Foundation and CF Canada; and the Natural Sciences and Engineering Research Council of Canada (RGPIN-2015-03742, 250053-2013). This work was also funded by the Government of Canada through Genome Canada (OGI-148) and supported by a grant from the Government of Ontario. The funders of the study played no role in study design, data collection and analysis, decision to publish, or preparation of the manuscript. S.K. is a trainee in the CANSSI Ontario STAGE (Strategic Training for Advanced Genetic Epidemiology) program at the University of Toronto.

